# BLINK: A Package for Next Level of Genome Wide Association Studies with Both Individuals and Markers in Millions

**DOI:** 10.1101/227249

**Authors:** Meng Huang, Xiaolei Liu, Yao Zhou, Ryan M. Summers, Zhiwu Zhang

**Affiliations:** Department of Crop and Soil Sciences, Washington State University, Pullman, Washington State, United States of America; Key Laboratory of Agricultural Animal Genetics, Breeding and Reproduction, Ministry of Education, College of Animal Science and Technology, Huazhong Agricultural University, Wuhan, Hubei, China; School of Electrical Engineering and Computer Science, Washington State University, Pullman, Washington State, United States of America

**Author notes:** Correspondence should be addressed to ZZ.

## Abstract

Big data, accumulated from biomedical and agronomic studies, provides the potential to identify genes controlling complex human diseases and agriculturally important traits through genome-wide association studies (GWAS). However, big data also leads to extreme computational challenges, especially when sophisticated statistical models are employed to simultaneously reduce false positives and false negatives. The newly developed Fixed and random model Circulating Probability Unification (FarmCPU) method uses a bin method under the assumption that Quantitative Trait Nucleotides (QTNs) are evenly distributed throughout the genome. The estimated QTNs are used to separate a mixed linear model into a computationally efficient fixed effect model (FEM) and a computationally expensive random effect model (REM), which are then used iteratively. To completely eliminate the computationally expensive REM, we replaced REM with FEM by using Bayesian information criteria. To eliminate the requirement that QTNs be evenly distributed throughout the genome, we replaced the bin method with linkage disequilibrium information. The new method is called Bayesian-information and Linkage-disequilibrium Iteratively Nested Keyway (BLINK). Both real and simulated data analyses demonstrated that BLINK improves statistical power compared to FarmCPU, in addition to a remarkable improvement in computing time. Now, a dataset with half million markers and one million individuals can be analyzed within five hours, compared with one week using FarmCPU.

## INTRODUCTION

Biomedical innovations have outpaced computing innovations since the completion of the human genome project^1, 2^. Genome-wide association studies (GWAS) have identified many genetic loci presumed to control some human diseases and agriculturally important traits^3–5^. However, a substantial proportion of these discoveries were false positives, attributed to a failure to consider population structure and cryptic relationships among individuals in the analyses^6–8^. Incorporating population structure and cryptic relationships as covariates dramatically reduces false positives, but also causes false negatives and computational burdens^9–11^.

Population structure is typically incorporated as a fixed effect in the General Linear Model (GLM), which is computationally efficient. Initially, population structure was derived as the proportions of individuals belonging sub-populations^12, 13^. Several alternatives for defining population structure, such as principal component analysis^14, 15^, were developed to further improve computational efficiency. Cryptic relationships can be incorporated in two ways. One way is to include all genetic markers as random effects. Some of these markers will capture the effects of Quantitative Trait Nucleotides (QTNs) through Linkage Disequilibrium (LD)^16–18^. The other way is to first derive kinship among individuals, using all genetic markers. The kinship is subsequently used to define the variance structure of individual effects as random effects. In the latter, both population structure and kinship can be incorporated into a fixed effect and random effect Mixed Linear Model (MLM)^9^. However, the computation of the MLM is intensive. Thus, multiple methods have been developed to reduce the computing times of MLM.

The first milestone that reduced this computational burden was the development of the Efficient Mixed Model Association (EMMA)^19^. Prior to EMMA methods, Maximum Likelihood (ML) or REstricted Maximum Likelihood (REML) functioned as the genetic variance and the residual variance. With ML or REML, optimization must be performed in two dimensions using methods such as Expectation and Maximization (EM). By using EMMA, ML or REML functions only as the ratio between genetic variance and residual variance. By reducing the optimization from two dimensions (genetic variance and residual variance) to one dimension (genetic-to-residual variance ratio), computing speed dramatically improves.

The second milestone was the use of empirical Bayesian estimation of population parameters such as genetic and residual variances or their ratio. This method is based on the assumption that each testing marker contributes only a small proportion of total genetic variance. Thus, population parameters for the testing markers can be approximated by the estimates from a reduced model without fitting each marker^20, 21^. Developed independently by two different groups, this algorithm has two names, Population Parameters Previously Determined (P3D**)^21^** and EMMA eXpedited (EMMAX)^20^.

Inspired by EMMA, EMMAx, and P3D, an exact algorithm, Genome-wide Efficient Mixed-Model Association (GEMMA)^22^, was developed to derive estimates of population parameters for each testing marker with the same computing speed as P3D and EMMAX.

The third milestone was the Compressed MLM (CMLM) that clusters individuals into groups based on kinship^21^. The computing time complexity of MLM is the cubic power of sample size. Clustering individuals into groups reduces the sample size from number of individuals to number of groups. Consequently, computing time is dramatically reduced in CMLM. Clustering individuals into groups is performed in a reduced model without fitting testing markers. The optimized grouping is used to test markers one at a time. The computing advantage of CMLM is greater for datasets with larger numbers of markers.

The fourth milestone was a method called Factored Spectrally Transformed Linear Mixed Models (FaST-LMM)^23^, which uses a rank-reduced kinship. Rather than using all available markers, a subset of genetic markers—less than the number of individuals in the sample—is used to create the rank-reduced kinship. Furthermore, FaST-LMM directly uses this subset of markers to define the relationship among individuals for ML or REML optimization without first calculating kinship. As a result, computing time is linear to subset size and independent of sample size.

The fifth milestone was GRAMMAR-Gamma^24^, a method that splits the association analysis into two steps. The first step uses MLM to derive the residual. The second step tests the residual as transformed traits in a fixed effect model and applies a correction factor to test statistical values. The computing complexity of the second step is completely linear to both number of individuals and markers.

With the exception of CMLM, the primary aim of the above milestones was to improve computing speed. The statistical power of each of these milestones remains similar to the conventional MLM^9^ because the same or similar kinship is used regardless of the traits being analyzed. CMLM, on the other hand, represented the first adjustment of kinship to improve statistical power^21^. In CMLM, genetic effects of individuals in the conventional MLM are replaced by the genetic effects of their corresponding kinship groups. That is, kinship among individuals is replaced by kinship among groups. Furthermore, the adjustment on kinship is optimized for the particular traits being studied. For example, the kinship with the best ML or REML is used for testing markers one at a time. Other optimizations were also developed to define minimum and maximum group kinship, in addition to average kinship^25^.

The second adjustment employs kinship that is not only specific for traits, but also specific for testing markers^26, 27^. Kinship is built by using only the markers that are associated with a trait. However, multiple associated markers can be genetically linked. To remove this redundancy, a bin procedure was developed within the Settlement of MLM Under Progressively Exclusive Relationship (SUPER) method to ensure that, at most, only one associated marker is selected from each bin^27^. Furthermore, the kinship changes according to testing markers to eliminate the confounding between kinship and testing markers. The trait-associated markers are excluded from the kinship calculation if they are also associated with the testing markers. This association is determined by linkage disequilibrium (LD) in SUPER^27^, or when the associated markers are on the same fragment (within 1Mb) as the testing markers^26^.

The third adjustment, known as the multi-locus mixed-model (MLMM) approach, applies weight to kinship^28^. In addition to random individual effects, this adjustment also fits multiple associated markers in the MLM to split the variance explained by kinship in a stepwise regression fashion. The forward stepwise regression stops when the kinship explains no variance. The multiple markers are then tested simultaneously without kinship.

Recently, a fourth adjustment, called the Fixed and random model Circulating Probability Unification (FarmCPU)^29^ was developed to completely eliminate the kinship in the model for testing markers. In FarmCPU, the marker-testing model becomes a Fixed Effect Model (FEM). To control spurious association for testing a marker, associated markers are fitted as covariates. To avoid the over-fitting problem, firstly, the whole genome is equally divided into certain number of bins, and only one significant maker with smallest P value is selected to be the candidates of covariates (pseudo QTNs). Secondly, these pseudo QTNs in the FEM are determined by a Random Effect Model (REM). The pseudo QTNs, from various bin with different width, are firstly ranked by P value, then, the best combinations between different bin and the number of possible QTNs are determined by REM. The two types of models (FEM and REM) are performed iteratively until no change occurs in the selection of pseudo QTNs.

Despite these valuable advancements, more innovative computing tools and analysis methods are needed. For example, although FarmCPU boosts statistical power in GWAS, its REM process remains computationally demanding. Additionally, the bin approach from SUPER requires that all QTNs be evenly distributed throughout the genome, which is hardly true. However, only one QTN can be selected as a covariate even if multiple QTNs are located in the same bin, which limits statistical power. Thus, a critical need still exists for a method that can increase both computing efficiency and statistical power.

## IDEA

Herein, we present a new statistical method that was inspired by this critical need and builds upon our previous method, FarmCPU. In the new method, we use 1) Bayesian Information Criteria (BIC) in a FEM to replace REML in the REM, and 2) linkage disequilibrium information to replace the bin method. As a result, we have completely eliminated the computationally expensive REM and the requirement that QTNs be evenly distributed throughout the genome (Fig. 1).

**Figure 1.**
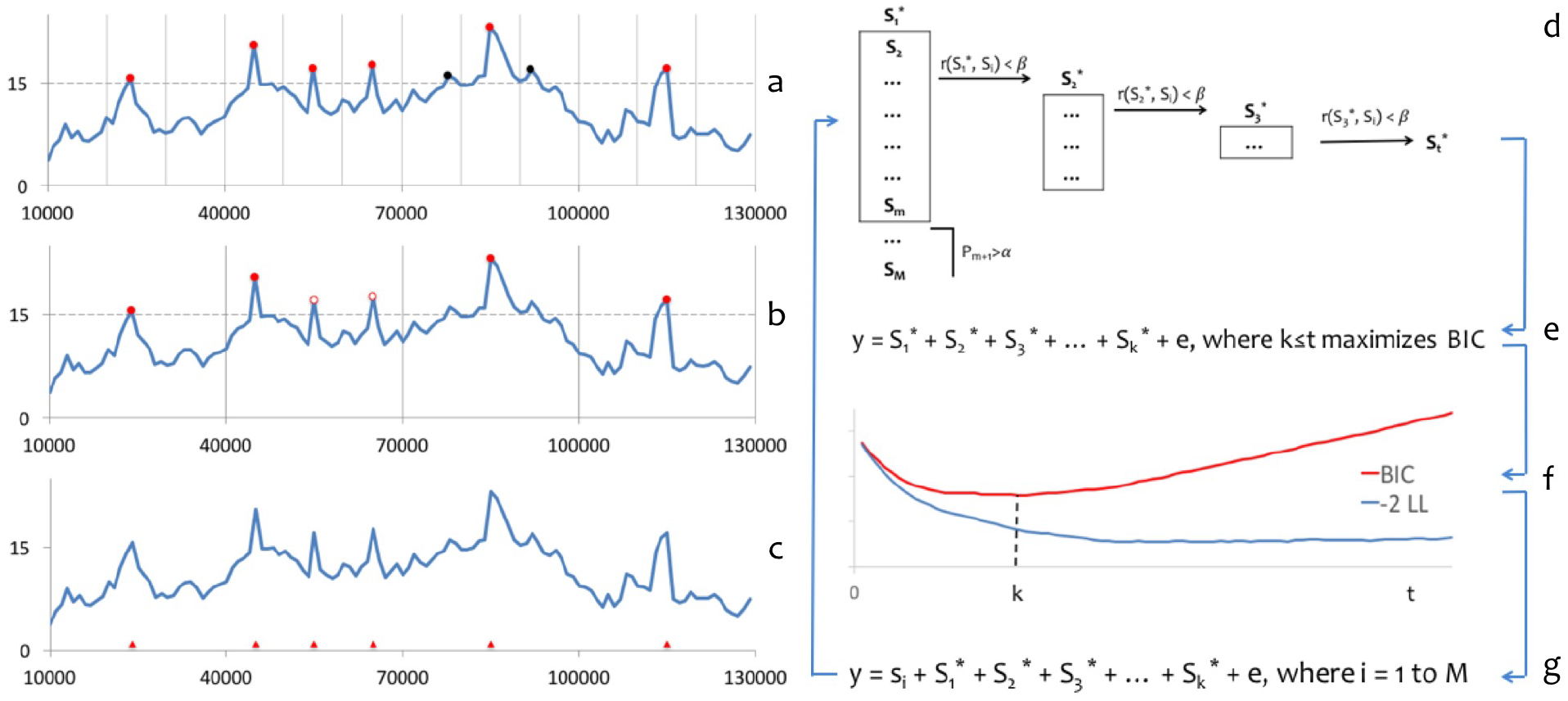
Limitation of the bin approach and proposed solution. Quantitative Trait Nucleotides (QTNs) are barely distributed evenly throughout the genome as required by the bin approach used in FarmCPU. The most significant marker from each bin, indicated by the filled red and black circles in **(a)** and **(b)**, is selected as a pseudo QTN if it passes a threshold (dash lines across vertical axes). A pseudo QTN could be false (filled black circle) if the bins (separated by the vertical lines) are too small **(a)**, or true, but not selected (open circle) if the bins are too big **(b)**, as illustrated by comparing the true QTNs (red triangles) positioned along the horizontal axis in **(c)**. Our alternative method is to sort the M SNPs first and filter them out if their P values are larger than a threshold (α). Among the m SNPs kept, additional SNPs are removed if their correlation (r) with the first SNP (S_1_^*^) is larger than a threshold (β). This process is repeated to select S_2_^*^, S_3_^*^,…, until the last SNP S_t_^*^ is selected **(d)**. As the t remaining SNPs are sorted, we fit the first k of them in a Fixed Effect Model (FEM) **(e)** and examine the corresponding twice negative log likelihood (-2LL) and Bayesian Information Criteria (BIC) **(f)**. As more SNPs are fitted, - 2LL continually improves (blue line), while BIC reverses (red line) because BIC applies a penalty with increasing numbers of SNPs. The k SNPs that give the best BIC are used as pseudo QTNs and fitted as covariates in another FEM to test all SNPs, one (s_i_) at a time, as described by the conceptual model **(g)**. This process (**d** to **g**) is iterated until the pseudo QTNs remain the same. We named this alternative solution the Bayesian-information and Linkage-disequilibrium Iteratively Nested Keyway (BLINK) method.

Our proposed method requires performing two FEMs in an iterative fashion. The first FEM tests genetic markers (SNPs), one at a time, and uses a set of associated markers as covariates. For the convenience of illustration, these associated markers are also named pseudo QTNs. The model can be written as:

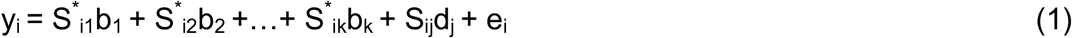

where y_i_ is the observation of the i^th^ individual; S^*^_i1_, S^*^_i2_, …, S^*^_ik_ are the genotypes of k pseudo QTNs, initiated as an empty set; b_1_, b_2_,…, b_k_ are the corresponding effects of the pseudo QTNs; S_ij_ is the genotype of the i^th^ individual and j^th^ testing marker; d_j_ is the corresponding effect of the j^th^ genetic marker; and e_i_ is the residual having a distribution with a mean of zero and variance of 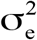.

Marker genotypes and P values from equation (1) are used to update the pseudo QTNs. Markers are filtered based on their P values by using a multiple test corrected threshold. The most significant marker is retained, while the markers in LD with the most significant markers are removed. Then, markers in LD with the second significant marker are removed, and so on, until no markers are in LD with each other. The first k of t remaining pseudo QTNs are fit into the second FEM, which is a reduced model that excludes the testing marker (S_ij_d_j_) compared with equation (1). The second FEM model can be written as:

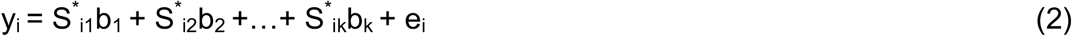

Bayesian Information Criteria (BIC) is examined with varied numbers of pseudo QTNs (k = 1 to t) until the optimum k is identified. We named the new method Bayesian-information and Linkage-disequilibrium Iteratively Nested Keyway (BLINK). The BLINK procedure is further detailed in the online methods section.

## RESULTS

We implemented the BLINK algorithm in two packages, one was written in R and the other in C. The R package was designed for the fact what more audiences are familiar with R than C. The C package was designed for computational efficiency. We named the two packages as BLINK-C and BLINK-R, respectively. As most of the analyses were conducted by BLINK-C, we simplified BLINK-C as BLINK. Both of these packages were compared with other two packages that are complementary for computing speed and statistical power. One is PLINK^30^ and the other is FarmCPU^29^. The package of PLINK was written in C and implemented the GLM method that has the minimum theoretical computing time complexity. The package of FarmCPU was written in R and implemented the FarmCPU algorithm that is superior to GLM in respect of statistical power. Statistical power was examined based on false positives, true positives, and statistical power at different levels of false discovery rate (FDR) and Type I error. To retain the real population structure (Fig. S1), we used phenotypes simulated from real genotypes that covered a wide range of species including human, one crop (maize), one livestock (pig), and two model species *(Arabidopsis* and mouse). Association study on real phenotype was conducted on flowering time in maize. Enrichment on a different study was performed to compare BLINK and FarmCPU. Finally, real genotype and phenotype data were also duplicated to synthetically create big data to examine observed computing time of BLINK-C, BLINK-R, PLINK and FarmCPU.

### Receiver operating characteristic

Using real genotypes from all five species, we simulated the QTNs controlling the phenotypes in two scenarios. The first scenario was a situation that rarely, if ever, occurs in practice—all QTNs were randomly located on the chromosomes without clusters. We call it “synthetic” scenario. The second scenario was a situation closer to reality—QTNs were clustered on chromosomes. Every two QTNs were restricted to next each other within 100 Kb. We call it “real” scenario. For each scenario, we examined statistical power under different levels of FDR and Type I error. FDR was defined as the proportion of false positives among the total number of positives identified. Type I error was derived from the empirical null P-value distribution of all non-QTN-bins. The relationship between statistical power and FDR or Type I error is described by the receiver operating characteristic (ROC) curves (Fig 2 and S3). The method with a larger area under curve (AUC) is preferred over the method with a smaller AUC. BLINK had a larger AUC than FarmCPU and PLINK for both power versus FDR and power versus Type I error; PLINK had a smaller AUC than FarmCPU and BLINK for both comparisons. This situation held true across all five species.

**Figure 2.**
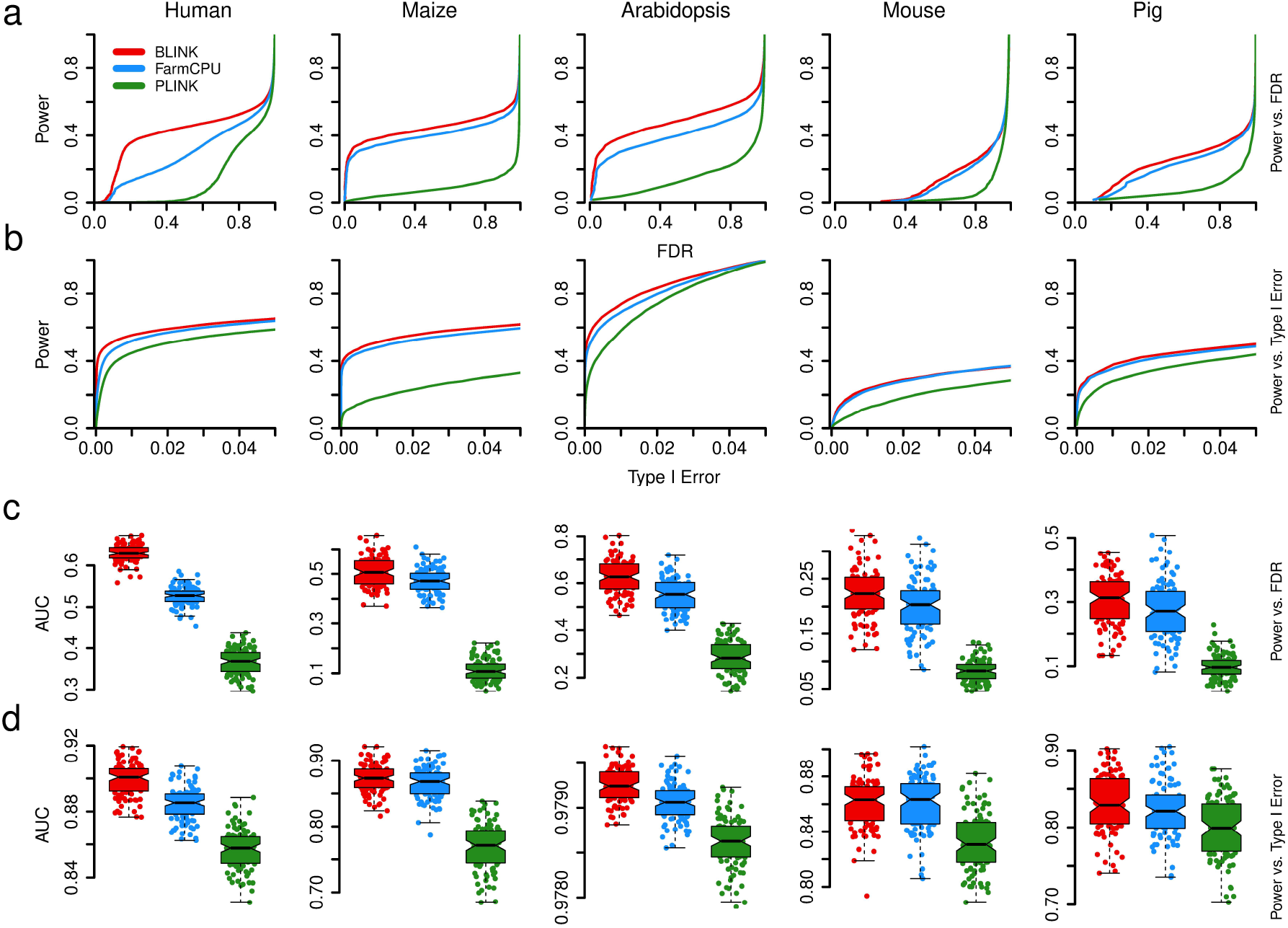
Statistical power and area under curve to detect clustered causal genes. Statistical power was defined as the proportion of simulated Quantitative Trait Nucleotides (QTNs) detected at cost defined by either False Positive Rate (**FDR**) or Type I error. The two types of Receiver Operating Characteristic (ROC) curves are displayed separately for FDR **(a)** and Type I error **(b)**. The Area Under the Curves (AUC) are also displayed separately for FDR **(c)** and versus Type I error **(d)**. Three GWAS methods (BLINK, FarmCPU, and PLINK) were compared with phenotypes simulated from real genotypes in five species (human, maize, *Arabidopsis thaliana*, mouse, and pig). The simulated phenotypes had a heritability of 75%, controlled by 500 QTNs for human, 100 QTNs for maize and mouse, and 50 QTNs for *Arabidopsis thaliana* and pig. These QTNs were randomly sampled from the available Single Nucleotide Polymorphism (SNPs) with restriction that every two QTNs were clustered within 300 Kb distance.

The model selection criteria (BIC) was compared with AIC (Akaike Information Criterion**)^31^** and EBIC (Extended BIC**)^32^** in all the five species examined. BIC overperformed other two model selection criteria (Fig S4). The determination of two markers in LD was based on the absolute values of their Person correlation coefficient. The default vales (0.7) was based on the comparisons of statistical power under different FDR (Fig S5).

### Associations and enrichment on real phenotype

We conducted GWAS on flowering time in maize using the three methods (BLINK, FarmCPU, and PLINK). PLINK exhibited strongly inflated P values (Fig. 3). For example, of the 397,323 SNPs in maize, PLINK identified 112,998 (28%) SNPs with P values smaller than the Bonferroni threshold. In contrast, neither FarmCPU nor BLINK demonstrated this problem. Results from BLINK and FarmCPU demonstrated that more than 99.9% of SNPs were not associated with flowering time after the adjustments by the associated SNPs.

**Figure 3.**
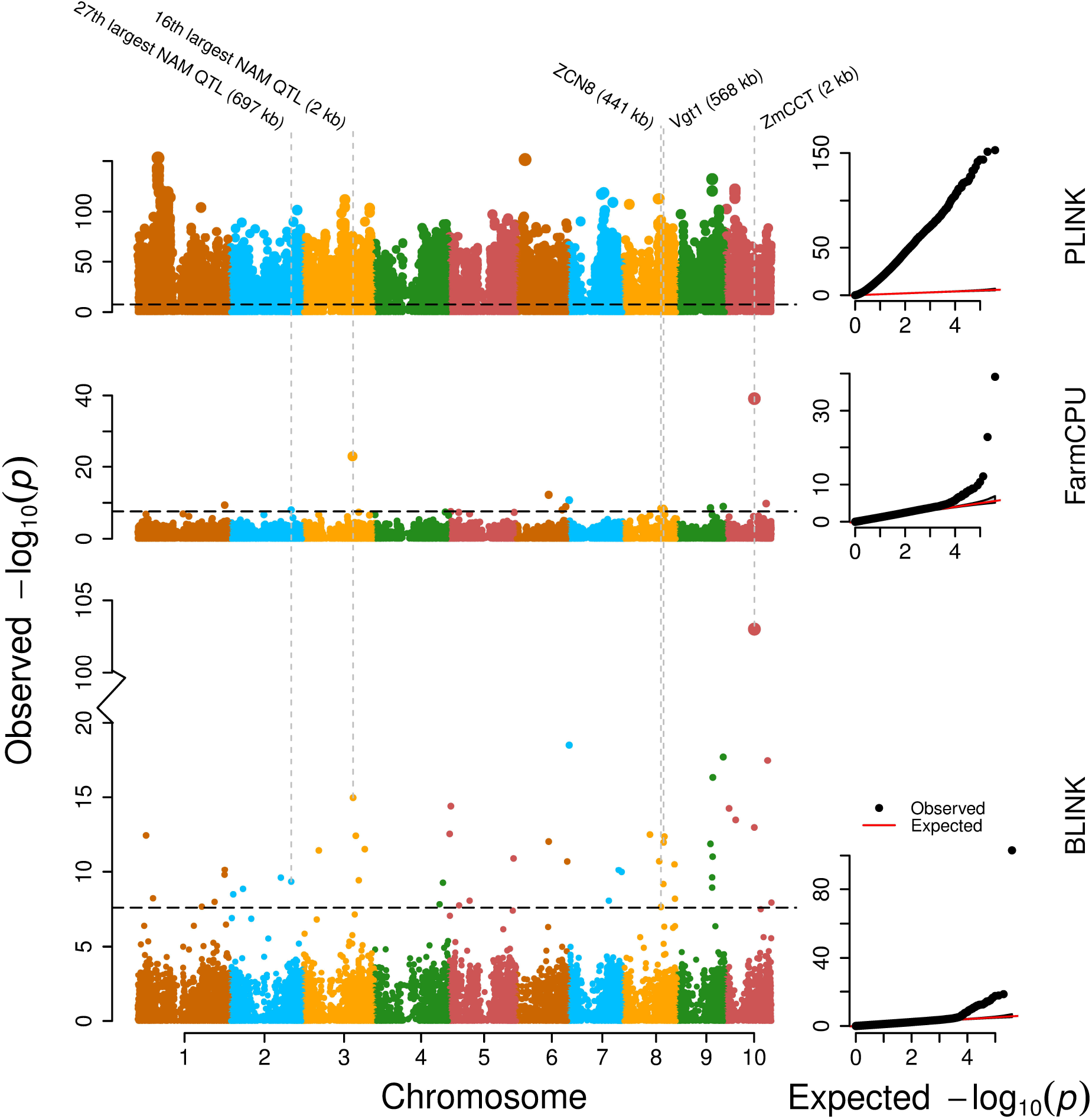
GWAS of flowering time (days to silking) in maize. The performance of three GWAS methods, BLINK, FarmCPU, and PLINK, are compared. The population included 2,648 individuals genotyped with 397,323 SNPs, after filtering out SNPs with a Minor Allele Frequency of 5% or less. All methods included the first two PCs and their products as covariates. The names of flowering-time candidate genes and NAM QTL that are surrounded by significant SNPs are labeled on the top, including the distance between significant SNPs and candidate genes/NAM QTL.

Notably, BLINK identified more associated SNPs than FarmCPU. FarmCPU identified 14 SNPs that passed the Bonferroni multiple test threshold (α = 0.01). In contrast, BLINK not only revealed 9 of these 14 SNPs, but also identified an additional 40 loci that passed the Bonferroni threshold. The significant SNPs identified by BLINK included the SNPs that are 2Kb from ZmCCT, 441kb from ZCN8, and 568 kb from Vgt1—three genes that have been previously cloned (Fig. 3). Compared to ZmCCT, which has a SNP at the same distance in both BLINK and FarmCPU, the distance between significant SNPs and the other two candidate genes are much closer in BLINK than in FarmCPU. Plus, BLINK identified three significant SNPs close to NAM QTL. BLINK and FarmCPU had much better control on inflated P values across the genome than PLINK. At the same level of control on inflated P values across the genome, BLINK identified more genetic loci associated flowering time.

Recently a distinct bigger population with 4,471 landraces was used to dissect genetic architecture of maize flowering time through GWAS. To distinguish the genes for the local environment adaptation, GWAS was conducted in controlled field experiments through a newly developed experimental design called F-one association mapping (FOAM). FOAM sampled individuals and cross them with small number of common parents to derive F1 families. GWAS on the evaluation of multiple trial F1 progeny identified 1003 genes associated with flowering time.

Both of genetic loci identified by FarmCPU and BLINK were significantly enriched on the 1003 flowering time genes identified by FOAM. The enrichment analyses defined flowering time genome regions as 50Kb upper stream and downstream of the 1003 genes. The FOAM flowering time gene regions occupied about 3% of maize genome. The strength of the enrichment was indicated by the contrast between the number of genetic loci hitting the FAOM flowering time gene regions observed and expected when the genetic loci were selected randomly. The derivations of the expected null distributions are illustrated in the Method section in details. The 9 common genetic loci identified by both FarmCPU and BLINK were enriched the most. Four out of nine genetic loci located in the flowering time gene regions. The chance to have four or above was below 1% by random chance. The five FarmCPU unique genetic loci were not enriched, but the 40 BLINK unique genetic loci were significantly enriched. Eight out of 40 located on the flowering time gene regions. The chance to have eight or above was below 5% by random chance. (Fig. 4).

**Figure 4.**
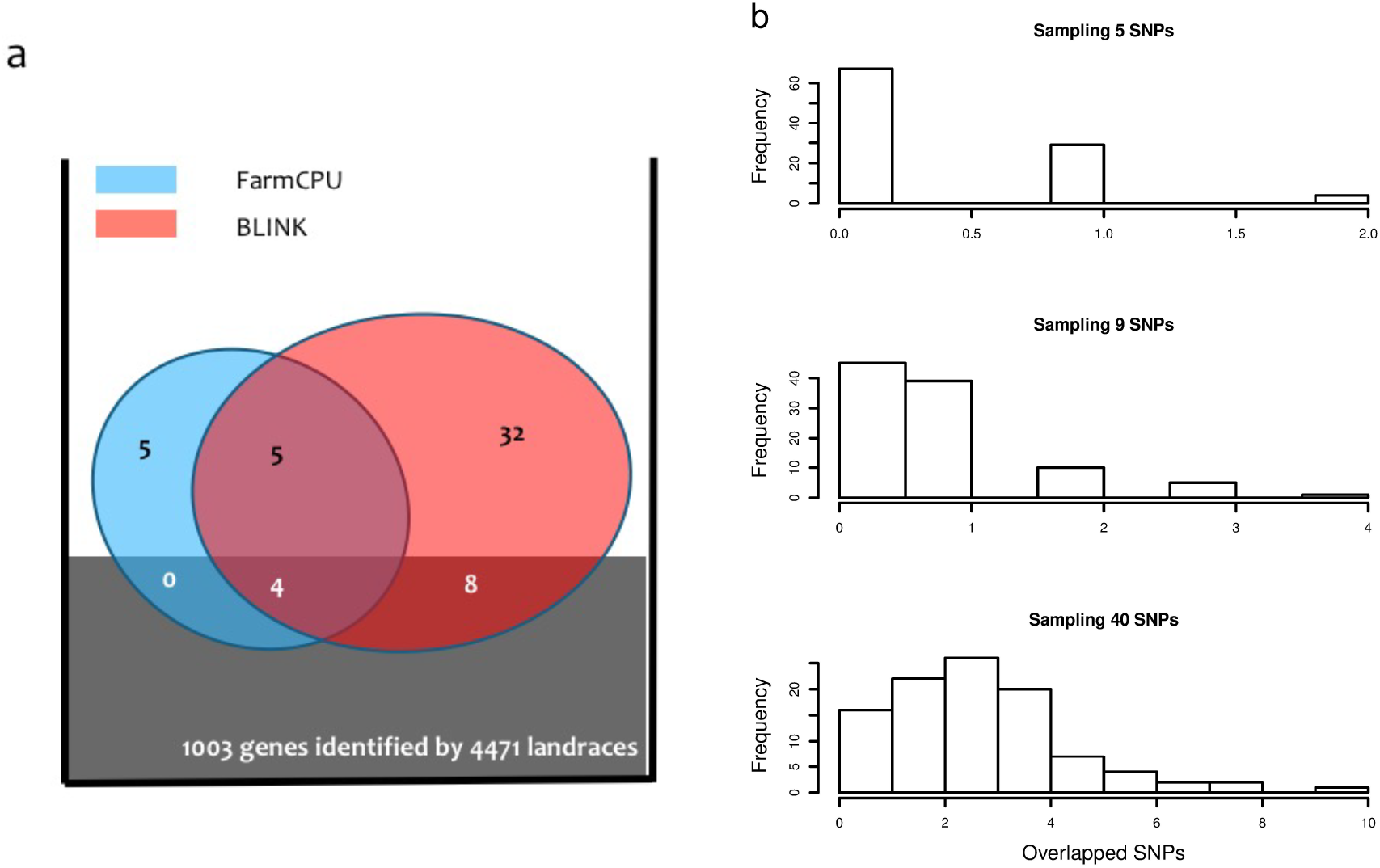
Enrichment of associated SNPs identified by BLINK and FarmCPU. The associated SNPs on maize flowering time were identified with Ames population containing 2279 lines by using BLINK and FarmCPU. These SNP were classified as the FarmCPU unique SNPs (5), common SNPs (9) and BLINK unique SNPs (40). The enrichment was performed as the overlap (within 50,000 base pair) with the 1003 flowering candidate genes identified by a separate population containing 4471 landraces **(a)**. The null probability distributions of are illustrated as the histograms of overlaps for randomly sampling five, nine and forty SNPs from the maize genome **(b)**. The FarmCPU unique SNPs were not enriched. The common SNPs and BLINK unique SNP were significantly enriched. The null probability was less than 1% for randomly sampling five SNPs with four or more overlapped with the 1003 candidate genes. Similarly, the null probability was less than 3% for randomly sampling forty SNPs with eight or more overlapped with the 1003 candidate genes.

### Theoretical computing time

In BLINK, association analysis of *M* markers, with *c* covariates, on a sample with *N* individuals takes a total computing time of c^2^MN. The quadratic term comes from the inverse of the left-hand side of the coefficient matrix. Both FarmCPU and BLINK add at most *t* pseudo QTNs as additional covariates to simultaneously control false positives and reduce false negatives. FarmCPU performs model selection of these pseudo QTNs with a REML procedure in the REM. The REM is solved to optimize bin size (b) and number of pseudo QTNs and to optimize the genetic-to-residual variance ratio with *p* iterations. FarmCPU’s computing time is tbp(c+t)^2^N for model selection and (c+t)^2^MN for association tests; total computing time is (M+tbp)(c+t)^2^N.

BLINK replaced REM with FEM for the model selection of pseudo QTNs. Consequently, the iterations are eliminated for optimizing the genetic-to-residual variance ratio. BLINK has a computing time of m^2^N, where m is number of markers with P values less than a threshold after a multiple test correction. The optimization of bin size is completely eliminated in BLINK. The total computing time for BLINK is (M+t)(c+t)^2^)N+m^2^n. The number of common covariates (c), pseudo QTNs (t), bin sizes (b), and iterations (p) are much smaller than both M and N. These scalars remain constant regardless of M and N sizes. Therefore, the computing time complexity is MN with respect to big O for all three methods, BLINK, FarmCPU, and PLINK (Table 1).

**Table 1.** Computing time complexity of BLINK compared with PLINK and FarmCPU*

| Methods | Model selection | Association Test | Total | Complexity over M and N |
| --- | --- | --- | --- | --- |
| **PLINK** | NA | $c^2MN$ | $c^2MN$ | $O(MN)$ |
| **FarmCPU** | $bsp(c+t)^2N$ | $(c+t)^2MN$ | $(M+bsp)(c+t)^2N$ | $O(MN)$ |
| **BLINK** | $t(c+t)^2N + m^2n$ | $(c+t)^2MN$ | $(M+t)(c+t)^2N + m^2n$ | $O(MN)$ |

\* The computing time is based on testing $M$ markers on a sample with $N$ individuals. All three methods contain common $c$ covariates. FarmCPU and BLINK add $t$ pseudo QTNs as additional covariates. FarmCPU examines $t$ QTNs over $b$ different levels of bin size and $s$ different levels of bin numbers. Using the EMMA algorithm, each examination optimizes the ratio of genetic-to-residual variance with $p$ iterations. BLINK selects $t$ pseudo QTNs with a computing time of $m^2n$ , where $n$ is the size of small subsamples of $N$ individuals. BLINK also eliminates optimization on bin size and on the genetic-to-residual variance ratio. The numbers of common covariates ( $c$ ), pseudo QTNs ( $t$ ), levels of bin size ( $b$ ), and iterations ( $p$ ) are much smaller than $M$ and $N$ . Therefore, the computing time complexity is $MN$ in respect of big O for all three methods.

### Observed computing time

The two BLINK software packages (C and R versions) gave the identical P values (Fig. S6). We compared their computing times with PLINK^30^ and FarmCPU^29^ for analyzing big datasets. The datasets were synthetically created by duplicating 8,800 human individuals with simulated phenotypes and with real genotypes of one-half million SNPs. The largest synthetic dataset contained about one million individuals. FarmCPU took about 11 hours to complete the analysis on a dataset with about 20 thousand individuals. During that same timeframe, BLINK-R completed the analysis on a dataset with about 100 thousand individuals. PLINK analyzed the largest dataset (one million individuals and one-half million SNPs) in about 10 hours. Amazingly, for the same dataset, BLINK-C completed the analysis in only 5 hours. Using a single-core CPU desktop computer, BLINK^®^ is about five times faster than FarmCPU, but 10 times slower than PLINK (new beta release testing version 1.90). BLINK-C is two times faster than PLINK, 20 times faster than BLINK-R, and 100 times faster than FarmCPU.

Among the four packages compared above, BLINK-C is the only software package that can fully use modern computer architecture with multiple cores for parallelization. We further examined the efficiency of BLINK-C on multiple-core computer systems. We tested BLINK-C on computers with core numbers ranging from 2 to 12 under different operating systems, including Linux and Mac. Results showed that the total computing time decreased linearly with number of cores (Fig. 5). For the dataset with about one million individuals and one-half million SNPs, a Mac Pro with 12 cores completed the analysis in just 30 minutes instead of 5 hours with a single core.

**Figure 5.**
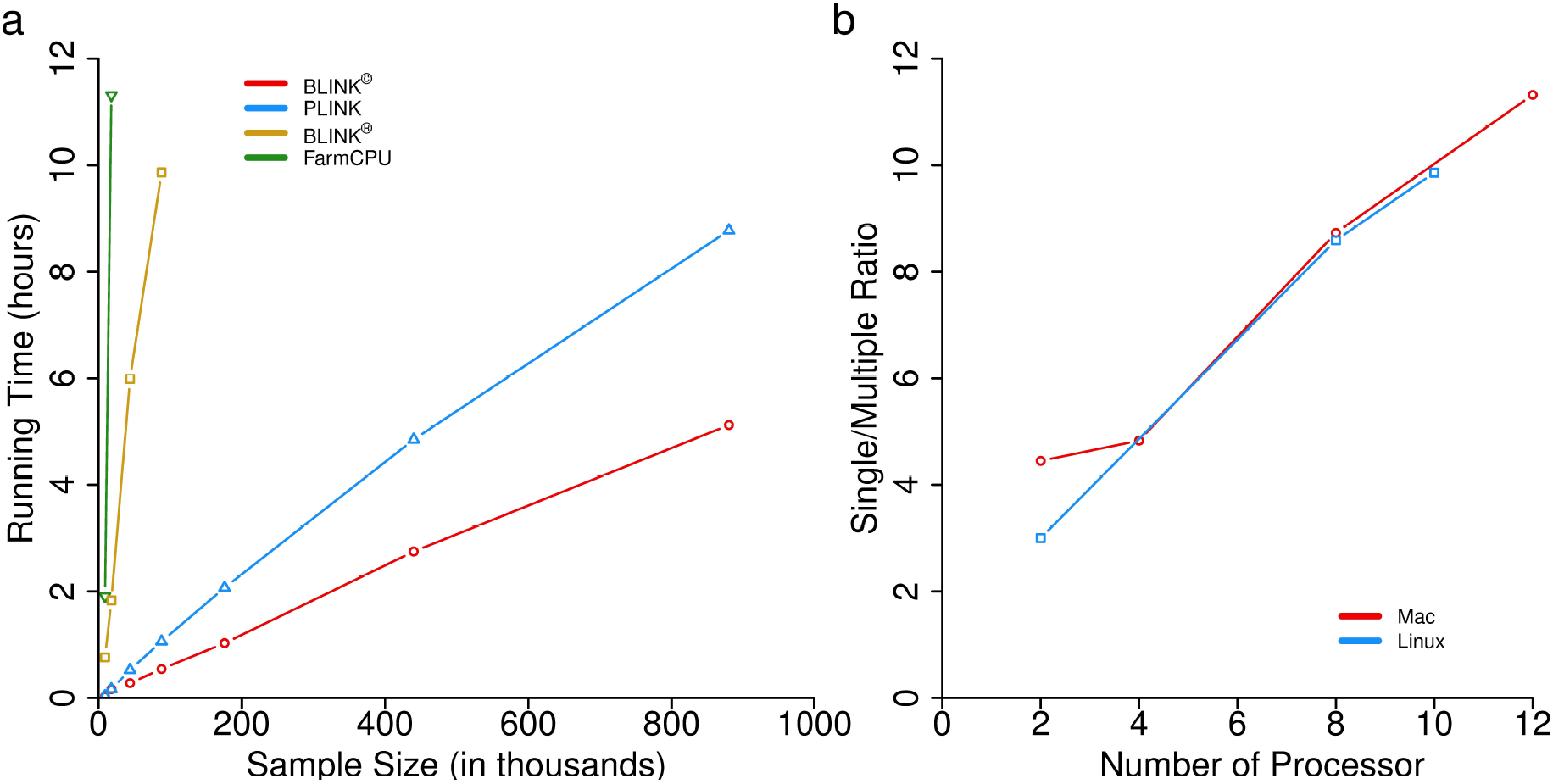
Comparison of computing time of BLINK and other software packages. The computing times using BLINK© and BLINK^®^ are compared with PLINK (version 1.90 beta) and FarmCPU **(a)** on synthetic dataset with duplication on original dataset containing 8,800 individuals genotyped with one-half million markers. Among the three software packages, BLINK© is the only one that can conduct parallel computation by using multiple central processing unit (CPU) cores. Different computers under different platforms were used to evaluate the efficiency of parallelization **(b)**. The efficiency is illustrated as the ratio of computing time of a single core to the computing time of two or more cores.

## DISCUSSION

Inspired by the critical need for computational efficiency and statistical power in big data analysis and by the most recently developed GWAS method, FarmCPU, we developed a faster and more powerful method. By substituting REML in FarmCPU’s REM with BIC in a FEM and by replacing the bin approach with LD, we achieved optimization in one dimension (number of pseudo QTNs) instead of two dimensions (number of pseudo QTNs and bin size). The optimization of the genetic-to-residual variance ratio was also eliminated by substituting REM with FEM, which directly solves residual variance without iterations. These improvements not only reduced computing time, but also simultaneously reduced false positives and false negatives.

### Substitution of REML with BIC

In both FarmCPU and BLINK models, markers are tested one at a time, with pseudo QTNs added as covariates to control false positives and reduce false negatives. FarmCPU selects these pseudo QTNs by using REM. Pseudo QTNs are used to derive kinship among individuals. The model chooses a set of pseudo QTNs to derive a kinship that provides the maximum likelihood^29^. Because FarmCPU does not gain extra parameters as more pseudo QTNs are included, the likelihood is not penalized for having more pseudo QTNs. In contrast, BLINK chooses pseudo QTNs using FEM. The more pseudo QTNs included, the greater the likelihood. Therefore, a penalty, such as BIC, on the number of parameters is necessary to identify the set of pseudo QTNs that best controls false positives and reduces false negatives. Both simulated data and real data demonstrated that BIC penalization worked well. By jointly using BIC and substituting for the bin approach, BLINK’s FEM performed even better than REM in FarmCPU.

### Robustness with genetic architecture

FarmCPU method uses bins as pseudo QTNs, according to the SUPER GWAS method^27, 29^. Both the number of bins (pseudo QTNs) and size of bins must be optimized together, in addition to optimizing the genetic-to-residual variance ratio. BLINK performs optimization in only one dimension (number of pseudo QTNs). A pseudo QTN represents a single SNP, not a bin. Multiple pseudo QTNs are acceptable regardless of proximity on the genome, unless they are in LD. In contrast, with FarmCPU, only one pseudo QTN can be selected if multiple pseudo QTNs are close enough to fall into the same bin. In practice, real QTNs are often clustered, rather than evenly distributed; thus, BLINK is more robust than other methods.

### Limitations

At the first iteration, the number of SNPs examined for LD is even more critical relative to both computing time and pseudo QTN selection. The P values may be severely inflated due to lack of control on false positives. Thus, the P threshold is much tighter at the first iteration compared to the rest of the iterations. For example, in the first iteration, the current default setting in BLINK© and BLINK^®^ only allows examination of SNPs with P values of 0.01 after a multiple test correction. For the remaining iterations, this P-value threshold is substantially released, 100% after a multiple test correction. When P values are extremely inflated at the first iteration, a stricter limit is set on number of markers (default is 1,000) examined for LD.

The selection of pseudo QTNs is significantly influenced by the threshold setting that determines whether a pair of SNPs are in LD. The current default setting used in BLINK© and BLINK^®^ is a Pearson correlation coefficient of 70%. Of course, the selection of pseudo QTNs is also determined by the optimization criteria, which is BIC in BLINK. We evaluated all of BLINK’s default settings in all five populations (Fig. S 4-5). Although these default settings work well, different criteria and/or methods may further improve optimization for specific species and/or datasets—a topic that remains open to future research.

Nevertheless, BLINK produced fewer false positives and identified more true positives than the most recently developed GWAS method, FarmCPU. BLINK outperformed FarmCPU^29^ and PLINK^30^ relative to both statistical power versus FDR and statistical power versus Type I error. The association analyses with BLINK identified many more genetic loci, including loci previously validated by other studies, than PLINK or FarmCPU. Although BLINK has the same computing time complexity as PLINK and FarmCPU, BLINK-C was not only faster than FarmCPU, but also faster than the most recently released beta testing version of PLINK. BLINK-C can analyze an extremely big dataset— one million individuals and one-half million markers—in 5 hours with a single core, or in 30 minutes with 12 cores. The software package BLINK-C has been released to the public as an executable program. The executable program of BLINK, help documents, demonstration data, and tutorials are freely available at http://zzlab.net/blink.

## MATERIALS AND METHODS

### BLINK Procedure

The BLINK method conducts two Fixed Effect Models (FEMs) and one filtering process that selects a set of pseudo QTNs, not in LD, as covariates. The entire sequence runs repeatedly until all genetic markers are tested and the selection of pseudo QTNs is optimized. The first FEM tests *M* genetic markers, one at a time. Pseudo QTNs are included as covariates to simultaneously control false positives and reduce false negatives. Specifically, the first FEM can be written as follows:

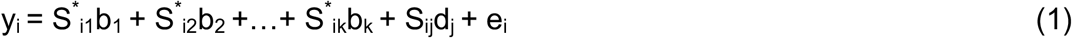

where y_i_ is the observation on the i^th^ individual; S_i1_, S_i2_,…, S_ik_ are the genotypes of k pseudo QTNs, initiated as an empty set; b_1_, b_2_, …, b_k_ are the corresponding effects of the pseudo QTNs; S_ij_ is the genotype of the i^th^ individual and j^th^ genetic marker; d_j_ is the corresponding effect of the j^th^ genetic marker; and e_i_ is the residual having a distribution with a mean of zero and a variance of 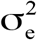. The primary goal of the first FEM is to calculate the P values for all *M* testing markers.

The second FEM is employed to optimize the selection of pseudo QTNs. Mathematically, the second FEM can be written as follow:

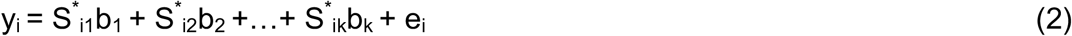

Equations (1) and (2) differ in two ways. First, the testing marker term in the first FEM is removed from the second FEM; therefore, no testing marker P values are output in (2). Second, the number of covariate pseudo QTNs are varied in the second FEM to select the optimum set of the first k out of t pseudo QTNs. The optimization is performed using Bayesian Information Criteria (BIC), which is twice the negative log likelihood plus the penalty on number of parameters, as follows:

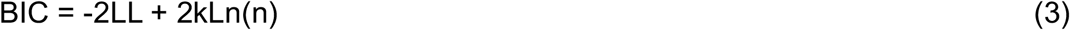

where LL is Log Likelihood, k is the number of pseudo QTNs, Ln is natural Log, and n is the number of individuals. There are t available pseudo QTNs that are sorted with the most significant at beginning and the lest significant at end. The first k pseudo QTNs are selected for examination with k varied from 1 to t.

All markers in equation (1) are candidates for pseudo QTNs in equation (2). These markers are filtered with two criteria: P value and LD. All markers are sorted first and then filtered out if their P values are larger than a threshold. Among the m SNPs kept, if their correlation (r) with the first SNP (S_1_^*^) is larger than a threshold, they are removed. This process is repeated to select S_2_^*^, S_3_^*^,…, until the last SNP, S_t_^*^, is selected (Fig 1).

As the t remaining markers are sorted and not in LD each other, the first set of k markers is more critical than the second set of k markers. We fit the first k markers in equation (2) and vary k until all possibilities are examined. The set of k markers with the best BIC is used as the set of pseudo QTNs in equation (1). This process is iterated until the pseudo QTNs remain the same. We named this alternative solution as Bayesian-information and Linkage-disequilibrium Iteratively Nested Keyway (BLINK) method.

### Genotype and phenotype Data

We used the exact same datasets we used in our previous publication for the FarmCPU method. These datasets covered five species including *Arabidopsis thaliana*, human, maize^35^, mouse^36^, and pig^37^. Markers with a Minor Allele Frequency (MAF) of 5% or below were filtered out from the original datasets. The number of individuals and markers and traits analyzed are summarized in Table S1. The principal components were calculated by PLINK using all the SNPs.

In maize genotype dataset, there are a total of 2,279 inbred lines, each line with 681,258 SNPs. The real phenotype of all the samples in this dataset is flowering time that was measured as days to silk (URL: http://www.panzea.org/#!genotypes/cctl).

The human dataset, “East Asian lung cancer dataset”, ID # phs000716.v1.p1, was obtained from dbGaP^5^. The name of this dataset is “East Asian lung cancer dataset” (ID # phs000716.v1.p1). Respecting the privacy and intentions of research participants, the dataset is only available under the permission of NIH (National Institutes of Health) and Intramural NCI (National Cancer Institute). There are a total of 8,807 individuals was involved into computing testing, each of them with 629,968 SNPs (URL: http://www.ncbi.nlm.nih.gov/projects/gap/cgi-bin/study.cgi?study_id=phs000716.v1.p1).

For the datasets of *Arabidopsis thaliana*, it contains a larger sample with 1,179 individuals that were genotyped with 214,545 SNPs (URL: http://archive.gramene.org/db/diversity/diversity_view).

### Synthetic data

The human dataset was synthetically duplicated to evaluate computing efficiency on large-scale datasets. The human dataset contained about one-half million (629,968) SNPs and 8,807 individuals. These individuals were duplicated 2, 5, 10, 20, 50, and 100 times. The largest synthetic dataset contained nearly one million (880,700) individuals. The number of SNPs remained the same, at approximately one-half million.

### Simulated phenotypes

The real genotypes of the five species were used to simulate phenotypes to examine statistical power under different levels of Type I error and FDR. The simulated phenotypes had heritability of 75%, controlled by a variable number of QTNs that were sampled from all real SNPs. Two scenarios were applied to the sampling of SNPs, without and with restriction. The restriction was that QTN must be within distance of 300 Kb with another QTN. The QTNs had effects that followed a normal distribution. These QTNs were summed together as the total additive genetic effect for each individual according to its real genotypes. Variance of additive genetic effect was calculated across all individuals. A normally distributed residual effect was assigned to each individual. The variance of the residual effect was assigned accordingly so that the proportion of additive genetic variance equaled heritability. Genomes were divided into 10KB-sized bins. Bins were classified as QTN bins if they contained a QTN, otherwise as non-QTN bins. The P value of a bin was represented by the most significant single nucleotide polymorphism (SNP). Statistical power was defined as the proportion of QTNs detected at different levels of false discovery rate (FDR) and Type I error derived from the empirical null distribution of non-QTN bins.

### Power, Type I error and FDR

Number of false and true positives were counted based on bins described in our previous study^29^. The size of bins was varied starting at a single base pair to one mega base pairs. We reported the results from using a 10 KB bin size. The P value of a bin was represented by the most significant SNP. A bin was considered a QTN-bin if it contained at least one QTN, otherwise, a non-QTN-bin. A non-QTN-bin with a P value that passed a threshold was counted as a false positive bin. A QTN-bin with a P value that passed the same threshold was counted as a true positive bin. The proportion of QTNs identified under different thresholds was calculated as statistical power. For all levels of statistical power, the proportion of non-QTN-bins was calculated as FDR. Type I error was derived from the empirical null distribution of all non-QTN bins. Furthermore, receiver operating characteristic (ROC) curves were used to compare statistical power under different levels of FDR and Type I error. The area under curve (AUC) was calculated with a starting point of zero and an ending point of one for FDR or Type I error.

## Author Contributions

ZZ conceptualized the study and wrote the manuscript. The concepts were implemented by MH in C language (BLINK-C), and by YZ in R language (BLINK-R). MH, XL, YZ and RMS performed the data analyses.

## Acknowledgements

This material is based upon work that is supported by an Emerging Research Issues Internal Competitive Grant from the Agricultural Research Center in the College of Agricultural, Human, and Natural Resource Sciences at Washington State University; Washington Grain Commission (Endowment and award number 126593); the National Institute of Food and Agriculture, U.S. Department of Agriculture (awards of 2015-05798 and 2016-68004-24770). The authors thank Dr. Linda R. Klein for valuable writing advice and editing the manuscript.

## Ethics Statement

Any opinions, findings, conclusions, or recommendations expressed in this publication are those of the authors and do not necessarily reflect the views of the funding agencies. All datasets analyzed herein were published previously. This study did not involve samples from humans or animals.

